# Speech-reception-threshold estimation via EEG-based continuous speech envelope reconstruction

**DOI:** 10.1101/2024.06.26.600765

**Authors:** Heidi B. Borges, Johannes Zaar, Emina Alickovic, Christian B. Christensen, Preben Kidmose

**Author notes:** Corresponding author: Preben Kidmose, Department of Electrical and Computer Engineering, Aarhus University, Center for Ear-EEG, Finlandsgade 22, 8200 Aarhus N, Denmark.

## Abstract

This study investigates the potential of speech-reception-threshold (SRT) estimation through electroencephalography (EEG) based envelope reconstruction techniques with continuous speech. Additionally, we investigate the influence of the stimuli’s signal-to-noise ratio (SNR) on the temporal response function (TRF). Twenty young normal-hearing participants listened to audiobook excerpts with varying background noise levels while EEG was recorded. A linear decoder was trained to reconstruct the speech envelope from the EEG data. The reconstruction accuracy was calculated as the Pearson’s correlation between the reconstructed and actual speech envelopes. An EEG SRT estimate (SRT_neuro_) was obtained as the midpoint of a sigmoid function fitted to the reconstruction accuracy versus SNR data points. Additionally, the TRF was estimated at each SNR level, followed by a statistical analysis to reveal significant effects of SNR levels on the latencies and amplitudes of the most prominent components. The SRT_neuro_ was within 3 dB of the behavioral SRT for all participants. The TRF analysis showed a significant latency decrease for N1 and P2 and a significant amplitude magnitude increase for N1 and P2 with increasing SNR. The results suggest that both envelope reconstruction accuracy and the TRF components are influenced by changes in SNR, indicating they may be linked to the same underlying neural process.

## 1 Introduction

Globally, an estimated 430 million people require hearing assistance, with projections suggesting this number will reach 711 million by 2050 (WHO, 2021). Hearing loss leads to communication difficulties and can have adverse effects on the individual, including social isolation, depression and cognitive decline (Li et al., 2014; Lin et al., 2013; WHO, 2021). While hearing aids can alleviate the challenges caused by hearing loss by providing amplification, speech intelligibility in noisy environments typically remains a challenge for hearing-impaired listeners with hearing aids. Measures of an individual’s speech-in-noise ability can thus be advantageous for better hearing-aid fitting, especially with regard to speech-enhancement processing (Zaar et al., 2024). Speech intelligibility in noise is quantified by the so-called speech reception threshold (SRT), which is a measure of the signal-to-noise ratio (SNR) at which 50% speech intelligibility is achieved. It is typically measured behaviorally, i.e., by presenting speech stimuli in background noise and asking the test participant to repeat back what they understood. However, this method has limitations, particularly its reliance on the participant’s ability and willingness to respond. Previous studies have shown promise in estimating the SRT using brain responses to matrix sentences recorded with electroencephalography (EEG) (Lesenfants et al., 2019; Vanthornhout et al., 2018). If the SRT can be accurately predicted from EEG, it implies that no active response is necessary. This could facilitate the measurement of SRTs, particularly in populations that cannot respond, such as infants and unconscious patients (Lesenfants et al., 2019; Van Hirtum et al., 2023a; Vanthornhout et al., 2018) and assist in evaluating the impact of hearing-aid algorithms.

Both linear decoding (backward-modeling) and encoding (forward-modeling) approaches have been used to predict speech intelligibility (Lesenfants et al., 2019; Vanthornhout et al., 2018). While the decoding model is a strong model, predicting one speech feature from multiple EEG channels (Alickovic et al., 2019), it lacks the ability to investigate the spatiotemporal pattern of the correlation (Haufe et al., 2014). The encoding model, by contrast, predicts the EEG channels from sound features (Alickovic et al., 2019) and thus yields a temporal response function (TRF) for each of the EEG channels. The TRF represents the “impulse response” that connects the sound feature at the input to the EEG channel signal at the output. Since the TRFs are from individual channels, the encoding approach allows for interpretation of the spatiotemporal pattern of brain activity (represented in the EEG signal) evoked by the stimulus feature. The TRF is akin to the event-related potential, but instead of having to repeat stimuli multiple times, the TRF allows to estimate how the brain responds to continuous stimuli.

The objective of this study is to investigate the feasibility of EEG-based SRT estimation. One concern when conducting audiological testing is whether the results are representative of the patient’s performance in their everyday sound environment. To address this, employing more naturalistic stimuli will enhance the ecological validity of the test. Therefore, instead of employing matrix sentences for speech testing and EEG measurements, as done in previous studies (Lesenfants et al., 2019; Vanthornhout et al., 2018), we use open-set sentences for speech testing and continuous speech for EEG measurements to enhance the ecological validity of the target speech stimuli.

Additionally, we aim to explore how changes in SNR affect the TRF in a young, normal-hearing population when using continuous speech. Previous research involving 5-year-olds has shown changes in the latency and amplitude of the P1 deflection of the TRF in response to changes in SNR of continuous speech in stationary speech-weighted noise (Van Hirtum et al., 2023b). As the N1 and P2 components are observed in the auditory evoked potential in the developed auditory system (Ponton et al., 2000), it is worthwhile investigating whether they manifest themselves in the TRF and how they vary across SNRs. Previous studies have reported that the latencies of TRF deflections decrease, and their amplitudes increase with higher SNR levels when using recordings from magnetoencephalography (MEG) (Ding and Simon, 2012). Building upon these studies, we hypothesize that changes in the SNR will induce changes in the TRF in young, normal-hearing listeners as well.

## 2 Material and methods

### 2.1 Participants

Inclusion of participants was based on the following inclusion criteria: native Danish speakers, right-handed, no neurological diseases and no dyslexia to a degree that significantly affects their everyday lives, aged between 18 and 30 years, and normal hearing. Inclusion was based on self-reporting, except for the normal hearing criterion, which was confirmed based on pure-tone audiometry. Normal hearing was defined as hearing thresholds of maximally 20 dB hearing level (HL) across frequencies ranging from 0.125 to 8 kHz (0.125, 0.25, 0.5, 0.75, 1, 1.5, 2, 3, 4, 6, 8 kHz), consistent with WHO guidelines for normal hearing (WHO, 2021). The hearing level was assessed in 5-dB steps. The participants received compensation in the form of gift cards and transport reimbursement for their participation. Prior to participation, all participants provided signed and informed consent. The study was approved by the Institutional Review Board at Aarhus University under approval number TECH-2022-004.

### 2.2 Experiments

The experiment was divided into three visits. During the first visit, the participants’ SRT was measured behaviorally (SRT_beh_), a reading span test (Daneman and Carpenter, 1980) and Edinburgh Handedness Inventory test (Oldfield, 1971) were conducted, and an ear-impression was made. The data recorded during the second visit is intended for another study and is therefore not further described here. During the third visit, the EEG experiment was conducted.

#### 2.2.1 EEG recordings

EEG data were recorded at a sampling rate of 4096 Hz using a Biosemi Active EEG (Amsterdam, Netherlands) setup with a 64-electrode EEG-cap and in addition two mastoid electrodes.

Additionally, 12 ear-EEG electrodes and one Fpz electrode were recorded with the SAGA32+/64+ system (TMSi, Oldenzaal, Netherlands). However, analysis of the ear-EEG data is not included in this study and has been deferred for future investigation. This article only presents results from the analysis of scalp electrodes.

#### 2.2.2 Stimuli presentation

In the EEG paradigm, the diotic stimuli were presented via a soundcard (RME Hammerfall DSB multiface II, Audio AG, Germany) and delivered through Etymotic ER-1 insert Earphones (Etymotic Research, Inc., IL, USA) connected to the ear-EEG earpieces via sound tubes. The experiment was conducted in a double-shelled RF shielded test box. The system was calibrated using an ear simulator type 4157 and the outer-ear simulator DB2012 (Brüel & Kjær, Denmark), with measuring amplifier type 2610/2636 (Brüel & Kjær, Denmark). During stimuli presentation, the speech was maintained at 65 dB SPL, while the noise level was varied to obtain the desired SNRs. The steady-state speech-shaped noise used as noise in both the behavioral SRT and EEG experiment was used for calibration.

The stimuli were presented in the same way in both the EEG experiment and the behavioral speech test, with the only difference that the stimuli were presented through disposable foam ear tips in the speech test and through the ear-EEG earpieces during the EEG recordings.

#### 2.2.3 Speech test

SRT_beh_ was assessed using the Danish Hearing-In-Noise Test (HINT) sentence lists (Nielsen and Dau, 2011), each consisting of 20 five-word meaningful sentences. The SNR at which 50% of the words were correctly repeated was chosen as the SRT_beh_ and determined using an adaptive procedure. In the adaptive procedure, the SNR for the next sentence was updated by adjusting the noise level by Δ*L* (in dB) relative to the noise level for the previous sentence. Δ*L* was calculated according to Δ*L* = (*p*_*previous*_ − *p*_*target*_) ∗ *g*, where *p*_*previous*_ is the proportion of words correct repeated in response to the previous sentence, *p*_*target*_ is the target proportion (here 0.5), and g is a constant, which was set at 8 dB for the first 4 sentences and at 4 dB for the remaining sentences (Bio-logic Systems Corp., 2005). The resulting changes in SNR depending on words correct repeated are shown in Table 1. This approach for adaptive SNR changes when using word scoring is similar to approaches described by Hagerman and Kinnefors (1995), Hernvig and Olsen (2005) and Brand and Kollmeier (2002). Two concatenated HINT sentence lists (i.e., 40 sentences) were used in the adaptive procedure, with the first sentence starting at a SNR of -10 dB. The SRT_beh_ was calculated as the mean SNR obtained with sentences 5-40, i.e., responses obtained with the smaller constant of g (4 dB).

**Table 1.** SNR changes depending on words correct.

| Words correct | SNR change [dB] for sentence 1-4 | SNR change [dB] for sentence 5-40 |
| --- | --- | --- |
| 0 | 4 | 2 |
| 1 | 2.4 | 1.2 |
| 2 | 0.8 | 0.4 |
| 3 | -0.8 | -0.4 |
| 4 | -2.4 | -1.2 |
| 5 | -4 | -2 |

*Note: SNR changes depending on the number of words correctly answered. Note that the step size is halved after the first 4 sentences.*

Initially, two concatenated HINT training sentence lists were introduced to the participant using an adaptive procedure. This was followed by two concatenated HINT sentence lists to adaptively determine the SRT_beh_. After the SRT_beh_ was determined, one HINT sentence list was run for each of 5 different SNR levels: SRT_beh_+4, SRT_beh_+2, SRT_beh_, SRT_beh_ -2 and SRT_beh_ -4 dB, as well as for a clean-speech condition, to obtain the words-correct score data. In the following, only the adaptively measured SRT_beh_ was used whereas the word-scoring data measured in the six mentioned conditions were not further processed.

#### 2.2.4 EEG-based SRT estimation

To estimate the SRT_beh_ from EEG (SRT_neuro_), audiobook excerpts were used as stimuli. Audiobooks provide naturally connected speech, offering better ecological validity than alternative options like concatenated HINT sentences. The audiobook excerpts of approximately 1 min duration, taking sentence boundaries into account, were presented to the participants at 5 different SNR levels relative to the individual participants’ SRT_beh_ (SRT_beh_+4, SRT_beh_+2, SRT_beh_, SRT_beh_-2, SRT_beh_-4 dB) and clean speech. In analogy to the behavioral speech test stimuli, the SNRs were obtained by using a fixed speech level of 65 dB SPL and adjusting the noise level. The audiobook material was filtered using a first-order lowpass filter with a cutoff frequency of 2 kHz to approximate the power spectral density of the HINT material. Fig. 1 illustrates the resulting mean third-octave band power spectral density for the audiobook, along with that of the HINT speech corpus. The power spectral density was extracted from each audio excerpt (approx. 1 min duration per excerpt) and from each HINT list by concatenating all HINT sentences belonging to the list (approx. 30 s duration per list).

**Figure 1.**
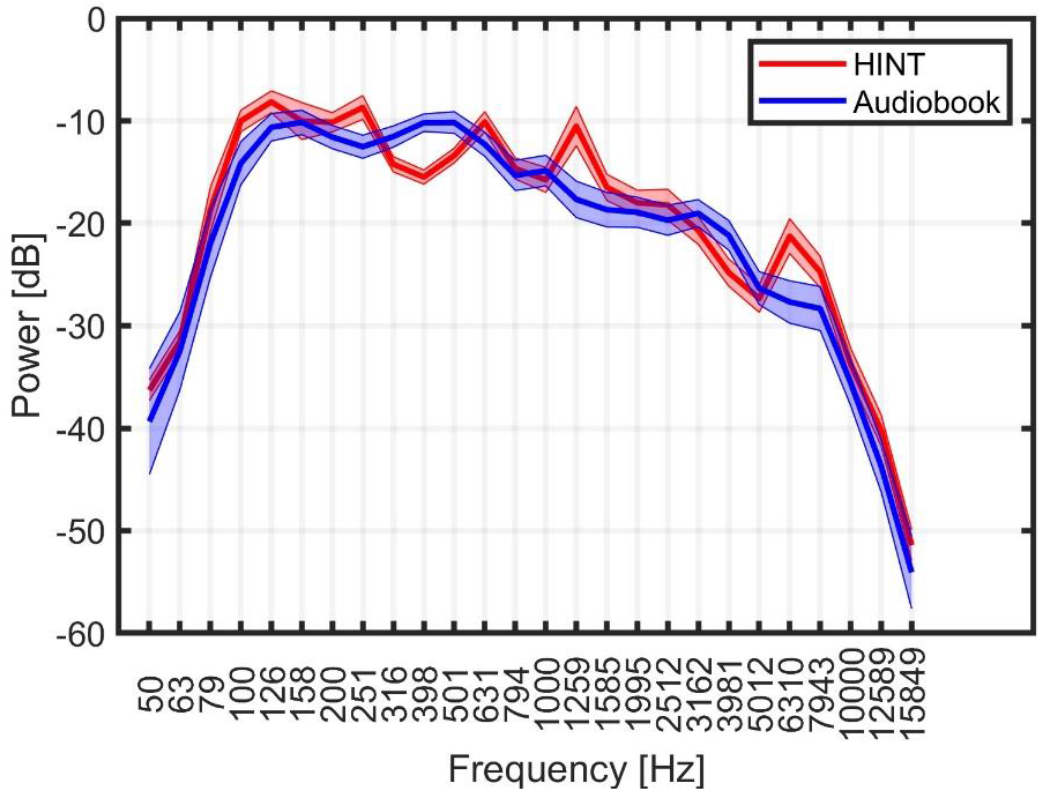
The one-third octave band long term spectral density for the HINT sentences and the audiobook material. The bold lines represent the mean spectral density, the shaded areas around them depict the range of ±2 standard deviations (SD) across the audio excerpts (audiobook material) or lists (HINT sentences).

The average fundamental frequency was 123 Hz for the audiobook material and 117 Hz for the HINT material. The noise started 2000 ms prior to the onset of the audiobook sound and ended 600 ms after the offset of the audiobook sound. The noise was faded in and out with a 400-ms raised-cosine ramp-on and ramp-off. The experiment consisted of two training trials to familiarize the participant with the task, followed by 6 blocks of 16 trials each. SNR conditions and clean speech were randomly presented in each block. After each trial, a content-related 2-AFC (two-alternative forced-choice) yes/no question was asked. A recreational activity was included between the blocks to help keep the participants engaged and attentive. This activity involved a small, unrelated task, such as describing one’s morning routine in detail.

### 2.3 Data analysis

#### 2.3.1 Stimuli preprocessing

All analysis was conducted using Matlab (The MathWorks Inc) and the mTRF Toolbox (Crosse et al., 2016). The audiobook stimuli were processed by extracting the broadband amplitude envelope as the absolute value of the analytical signal obtained via the Hilbert transform. A power law was applied according to *x*(*t*)^0.6^, where x is the envelope and t is time, to compensate for the compression in the auditory system (Biesmans et al., 2017). The signal was then bandpass filtered between 1 and 8 Hz using 3^rd^-order high- and lowpass Butterworth filters, which were employed forward and backward using Matlab’s “filtfilt” function, overall resulting in a 12^th^-order bandpass filter, and finally down-sampled to a sampling rate of 64 Hz.

#### 2.3.2 EEG preprocessing

The EEG data were processed in two separate ways: one to optimize the independent component analysis (ICA) weights (the “ICA pipeline”) and another to process the EEG data for the decoder (the “EEG pipeline”). To obtain the ICA weights, the pipeline involved down-sampling the data to 256 Hz to reduce processing time. This was followed by bandpass filtering between 1 and 100 Hz using 3^rd^-order high- and lowpass Butterworth filters, again employed forward and backward using Matlab’s “filtfilt” function, resulting in a 12^th^-order bandpass filter. Subsequently, a Zapline Notch filter was applied, as described by de Cheveigné (2020). Channels with an RMS-value larger than 3 times the mean RMS-value across all channels were labelled as bad channels and removed. The removed channels were replaced using spherical interpolation. The channels were re-referenced to their common average, and all trials were epoched and concatenated. Finally, the “runica” function from eeglab (Delorme and Makeig, 2004) was applied to obtain ICA weights.

The EEG pipeline was applied by first down-sampling the EEG data to 256 Hz, removing the channels labelled as bad in the ICA pipeline, and reconstructing them using spherical interpolation. The EEG signal was then re-referenced to the average of all channels. Next, the ICA weights were applied to the data and any components labelled as ≥75% likely to be artefacts from eyes, heart, line noise or channel noise by the IClabel algorithm (Pion-Tonachini et al., 2019), were removed.

Following this, the data were bandpass filtered between 1 and 8 Hz using 3^rd^-order high- and lowpass Butterworth filters, which were employed forward and backward using Matlab’s filtfilt function. Then the data were resampled to 64 Hz and normalized by dividing each channel with the mean of the standard deviation (SD) across all channels. Finally, the channels were referenced to the average of all channels.

#### 2.3.3 SRT estimation

A decoder was used to estimate the SRT_neuro_, with training and testing conducted at the participant level. The decoder was trained on 16 trials with clean (no noise) speech, and subsequently applied to the conditions with noise. The reconstruction accuracy was calculated as the Pearson’s correlation between the reconstructed envelope and the actual envelope of the speech stimulus. To train the decoder, a decision window of *τ* = [−100 350] ms was used and the regularization parameter, λ, were optimized in the range [10^−4^ − 10^10^]. One λ value per participant was used for decoder training. The participant’s specific λ value was determined by selecting the value yielding the greatest reconstruction accuracy when performing a leave-one-out cross validation of all the trials in the clean speech condition. Reconstruction accuracies for EEG data measured in the noise conditions were obtained using a decoder trained on all trials of the clean speech condition. A one sided permutation test was performed with a precision of 0.01 and α=0.05 to determine whether the reconstruction accuracy increased with increasing SNR at the population level (in general for all participants). To that end, the following conditions were tested against each other: SRT_beh_-4 dB and SRT_beh_-2 dB; SRT_beh_-2 dB and SRT_beh_; SRT_beh_ and SRT_beh_+2 dB; SRT_beh_+2 dB and SRT_beh_+4 dB.

The noise floor was determined by using mismatched envelopes from 68 audio excerpts. These excerpts were extracted from the same audiobook as the stimuli presented to the participants, underwent the same preprocessing, and were of approximately the same length. However, they were not part of the speech stimuli presented to the participants. Reconstruction accuracy was then calculated as the Pearson’s correlation between the reconstructed envelope from each trial with noise (80 trials) and the 68 mismatched envelopes. To ensure equal length of the two envelopes, the EEG used for envelope prediction, or the mismatched envelope was cut to the length of the shortest duration of the two. Subsequently, the mean reconstruction accuracy was calculated to obtain one noise floor value per participant. To find a reconstruction accuracy for the clean speech condition, leave-one-out training was used on trials only from that condition.

A sigmoid function described in (Farris-Trimble and McMurray, 2013) was fitted to the reconstruction accuracy-vs-SNR data from single participants. The sigmoid function is parameterized as:

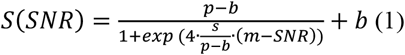

where *p* is the mean reconstruction accuracy of the clean speech condition, *b* is the reconstruction-accuracy noise floor, *s* is the slope and *m* is the midpoint-value, the midpoint value is used as the SRT_neuro_. We hypothesize that this is a potentially biased estimator of SRT_beh_, since reconstruction accuracy and % of words correct are not the same parameter. Thus, implementation of an offset from the midpoint of the fitted sigmoid function might be necessary to estimate SRT_beh_. During the fitting procedure, boundaries of [-10^10^, 10^10^] were imposed for the m value, and [0, 10^10^] for the *s* value. To ensure that the fit was conducted on an increasing trend, fits was only performed on participants where the clean speech condition was significantly higher than the noise floor. This was tested using a permutation test with 5% significance level. Each fitting was then repeated 100 times with a random initialization point for the *m* value ±10 dB from -2.52 dB, the average SRT_beh_ value measured for normal-hearing listeners found in the validation of the Danish HINT test (Nielsen and Dau, 2011). This SRT_beh_ was based on sentence-based scoring, since no word-based scoring was reported. Subsequently, the average of the sigmoid parameters for the 10 fits with the highest *R*^2^ values was chosen as the final fit. This procedure was used to mitigate issues related to local minima and enhance robustness of the fitting method. An example of a fit using data from a representative participant is shown in Fig. 2. The entire described fitting procedure was repeated 100 times to assess the test-retest reliability of the method.

**Figure 2.**
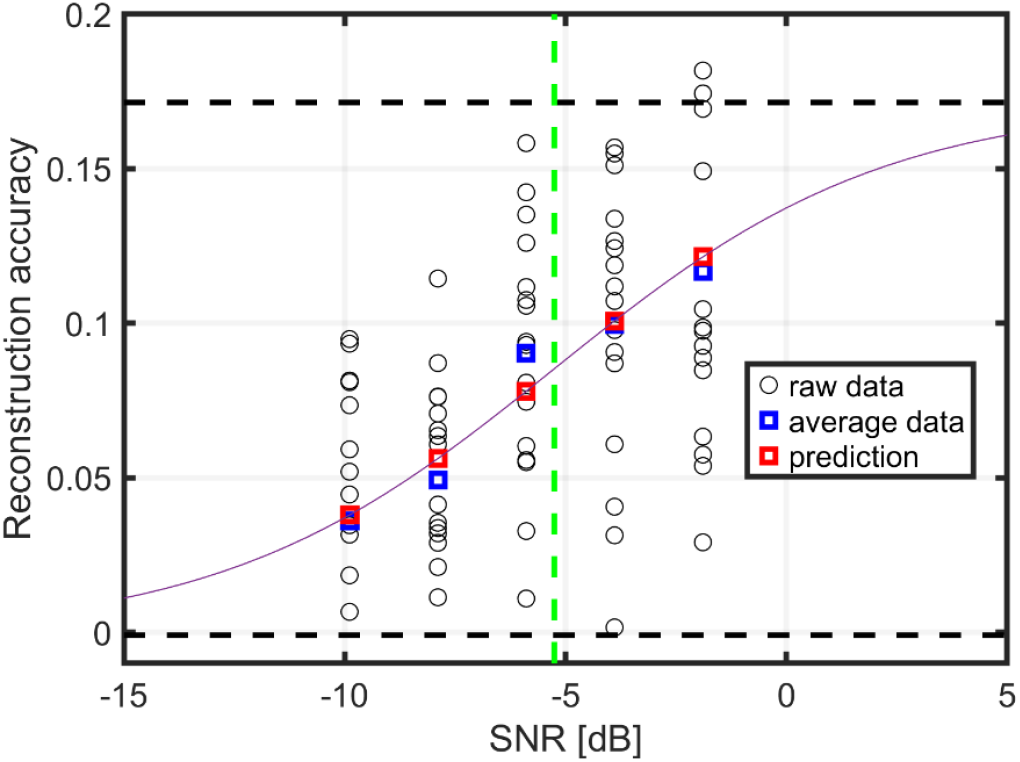
Example of a sigmoid fit for the reconstruction accuracy-vs-SNR data for one participant. Each circle represents the reconstruction accuracy for a single trial, the blue squares represent the average reconstruction accuracy for each SNR condition, the red squares show the datapoints predicted by the function for the considered SNR conditions. The dashed black lines represent the p and b coefficients of the sigmoid fit and the green dashed line the m coefficient of the sigmoid fit used for the SRT_neuro_ determination.

#### 2.3.4 Temporal response function

To compute the TRF, an encoder was trained using leave-one-out training within each condition. A hyperparameter (λ) value of 100 was used for all participants, as it yielded the highest mean prediction accuracy when testing the hyperparameters in the range [10^−6^ − 10^5^]. The λ was fixed not only across conditions but also across participants to facilitate the comparison of TRFs across conditions and participants. The time window investigated, *τ* = [−100 500] ms, was chosen to include the major components of the TRF. Model performance was assessed by calculating Pearson’s correlation between the predicted and the true EEG signal, termed prediction accuracy. A global selection of 17 electrodes was chosen based on high prediction accuracies, mirroring an approach used in a previous study (Fuglsang et al., 2017). A permutation test was performed to assess whether the prediction accuracy of the 17 electrodes in each condition, on a participant level, significantly exceeded that of the noise floor (α=0.05). The noise floor was calculated using the encoder model and mismatched envelopes, akin to the approach used in the decoding analysis. Only the TRFs with significant prediction accuracies were included in the further analysis. The mean TRF was calculated from the 17 chosen electrodes for each participant and SNR. Then the latencies and amplitudes of the two positive deflections (P1 and P2) and the one negative deflection (N1) were found by fitting a gaussian function: 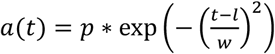, where *a* is amplitude, *p* is the deflection, *t* is time, *l* is the latency of the deflection and *w* is the width of the deflection. The boundary parameters used for the fit are shown in Table 2.

**Table 2.** Boundary parameters.

| Deflection | p[a.u.] | l[ms] | w[ms] |
| --- | --- | --- | --- |
| **P1** | [0 5] | [0 100] | [0 50] |
| **N1** | [-5 0] | [70 150] | [0 50] |
| **P2** | [0 5] | [100 300] | [0 50] |

*Note: Boundary parameters for gaussian function fit on the TRFs.*

The gaussian function fit was excluded from further analysis if it obtained an *R*^2^ value of less than 0.5. The relationship between the SNR, latency, and amplitude for the different deflections was assessed using a linear mixed model in RStudio (Posit team, Boston, MA, version 2023.12.1.402) with the nlme package (Lindstrom and Bates, 1990; Pinheiro and M. Bates, 1996) and maximum likelihood criteria. Participant number was modeled as a random effect (P), allowing for an individual offset for each participant. The latency and amplitude features (F) of the deflections were estimated using the model: F ∼ SNR + (P|1). Residuals were analyzed for normality by observing the qq-plot and histogram of residuals. Statistical significance of the fixed effect was assessed with a univariate Wald test with α=0.05. A Holm-Bonferroni correction was applied to avoid familywise errors, as six significance tests were performed, one for each feature. The clean speech condition was not included in the statistical test.

## 3 Results

### 3.1 SRT estimation

Twenty-two normal hearing participants were recruited for this study. However, due to technical problems, data recordings from two participants were incomplete, leading to their exclusion from the analysis. Consequently, the final analysis included data from 20 participants, comprising 17 females and 3 males, aged 18-29 years (mean 24.3 years). The highest pure-tone threshold across the two ears is shown as a function of audiometric frequency in Fig 3A. The SRT_beh_ ranged from - 6.00 to -3.28 dB with a mean of -5.35 dB and are shown in Fig 3B.

**Figure 3.**
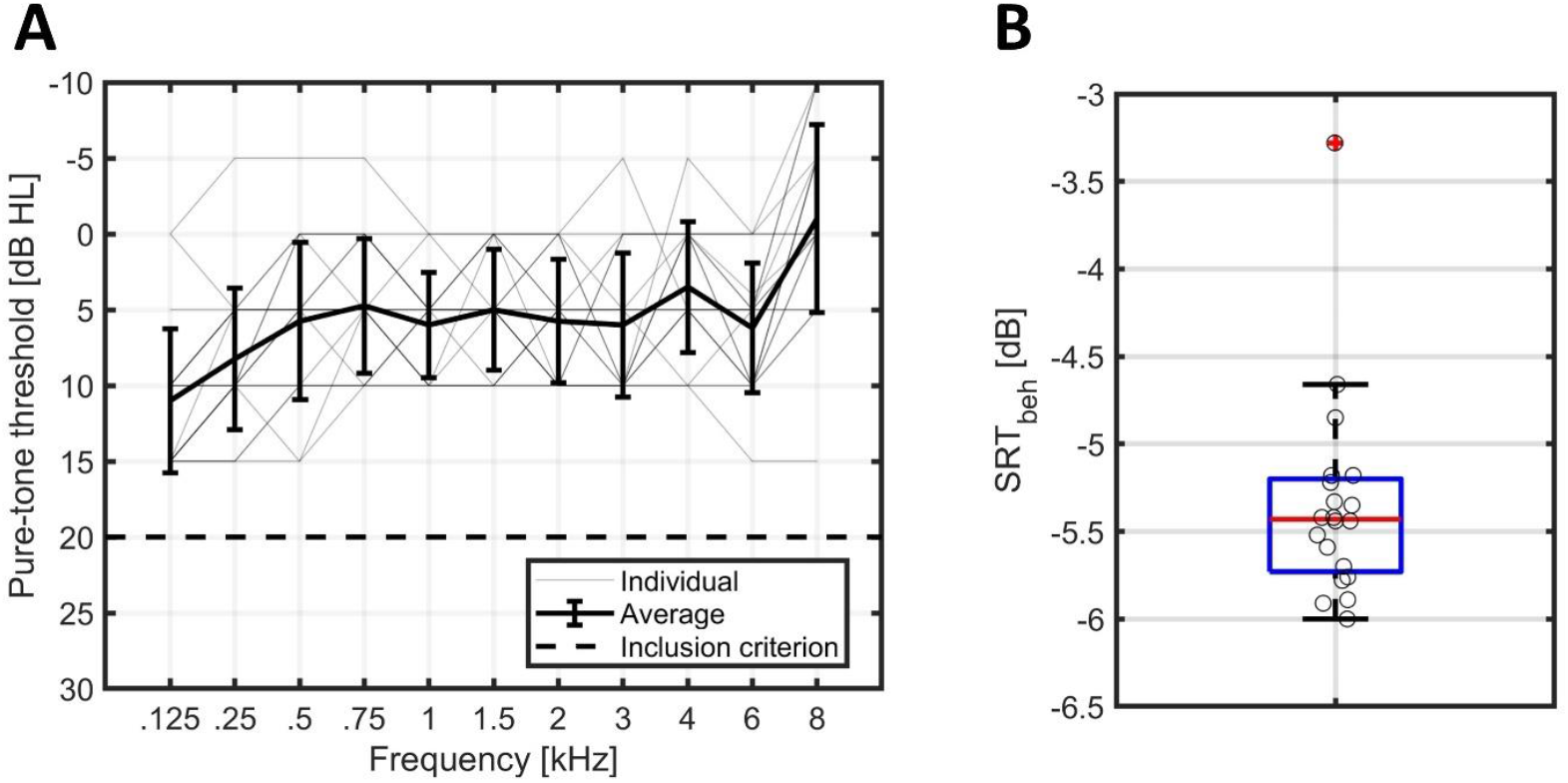
A: Highest pure-tone threshold across ears for each of the participants (thin lines), along with the mean and standard deviation across participants (shown in bold lines). The inclusion criteria of max 20 dB hearing level is shown as a dashed line. B: Boxplot of the measured SRT_beh_. The red line shows the median, the edges of the blue box the 25th and 75th percentiles, the black whiskers mark the most extreme values for non-outliers, and the red cross marks datapoints considered outliers, defined as a value more than 1.5 times the interquartile range away from the bottom or top of the box. Data for individual participants are shown as circles and are jittered slightly along the x-axis to improve the readability of the figure.

To investigate the feasibility of estimating the SRT from EEG measurements, we analyzed the reconstruction accuracy as a function of the SNR. Fig. 4A shows the reconstruction accuracy as a function of SNR (relative to the behaviorally measured SRT) for each participant (thin lines) as well as the mean across participants (bold line). The mean reconstruction accuracy exhibits a monotonic increase. Permutation tests showed a significant increase in the reconstruction accuracy for each 2 dB increase in SNR (all p-values below 10^−3^).

**Figure 4.**
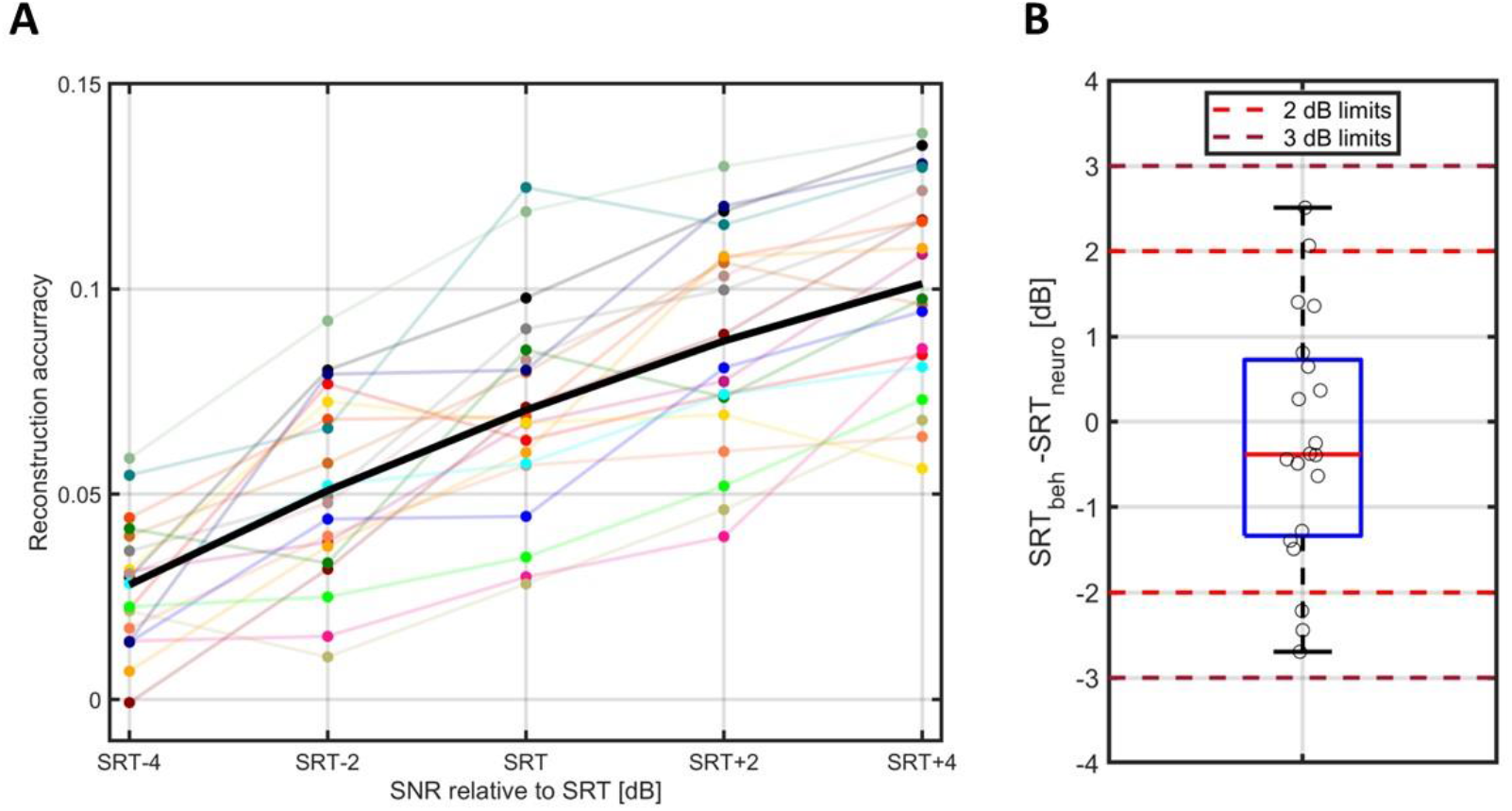
A: Reconstruction accuracy as a function of SNR (relative to the individual participant’s specific SRT_beh_) for each individual (dots, connected by thin lines) and the mean across participants (bold line). B: Boxplot of the difference between the SRT_neuro_ and SRT_beh_. The solid red line shows the median, the edges of the blue box the 25th and 75th percentiles and the black whiskers mark the most extreme values. Data for individual participants are shown as black circles and are jittered along the x-axis. The light red and dark red dashed lines indicate an absolute difference of 2 and 3 dB, respectively.

The individual SRT_beh_ (shown in Fig. 3B) was used to determine the SNR for the audiobook stimuli employed in the EEG recordings. With the SRT_beh_ ranging from -6.00 to -3.28 dB, the corresponding SNRs for the individual participants spanned from -6.00±4 dB to -3.28±4 dB in 2 dB steps (SNRs of -10 to 0.72 dB). The SRT_neuro_ was estimated as the midpoint-value from the fitting of the sigmoid function to the individual reconstruction accuracy for each participant The difference between the SRT_neuro_ and the SRT_beh_ is shown in Fig. 4B. The median of the individual difference between SRT_neuro_ and SRT_beh_ was very close to zero (0.38 dB), with group-level median values of - 5.43 dB for SRT_beh_ and -5.17 dB for SRT_neuro_. The standard deviation of the difference was 1.45 dB. We found that 15 out of the 20 participants had a difference between SRT_neuro_ and SRT_beh_ within ±2 dB and all participants had a difference between SRT_neuro_ and SRT_beh_ within ±3 dB. The maximum test-retest difference of the fitting procedure within participants was 1.42 dB.

### 3.2 Temporal response function

The grand average TRF exhibited three distinct deflections within the time window of 0 to 300 ms, consisting of two peaks around 50 ms and 200 ms (P1 and P2) and a trough around 120 ms (N1). The grand average TRF is shown in Fig. 5A for each of the 5 SNR conditions and the clean speech condition. TRFs for individual participants are provided in Appendix A1, note that the clean speech condition was not included in the statistical analysis of the TRFs. The prediction accuracy values from the encoding model were higher centrofrontally compared to the rest of the scalp, increasing with increasing SNR (Fig. 5B). The *p*-values obtained from the Wald test and the mixed-model values for the fixed effects are presented in Table 3. Statistically significant values were observed for the latency of N1 and P2 as well as the amplitude of N1 and P2. The negative coefficient of the latency fixed effect of the components P2 (-6.001) and N1 (-1.517) indicates that the latencies decreased with increasing SNR. The amplitude coefficients of the significant components N1(-0.015) and P2 (0.018) indicates that the magnitude of the components amplitude increased with higher SNR. Fig. 6 shows the latency, and amplitude estimates for the three deflections (denoted P1, N1 and P2) of the TRF across the six conditions. From Fig. 6 we observe a general pattern of decreasing latencies and enhanced amplitude with increasing SNR.

**Table 3.**
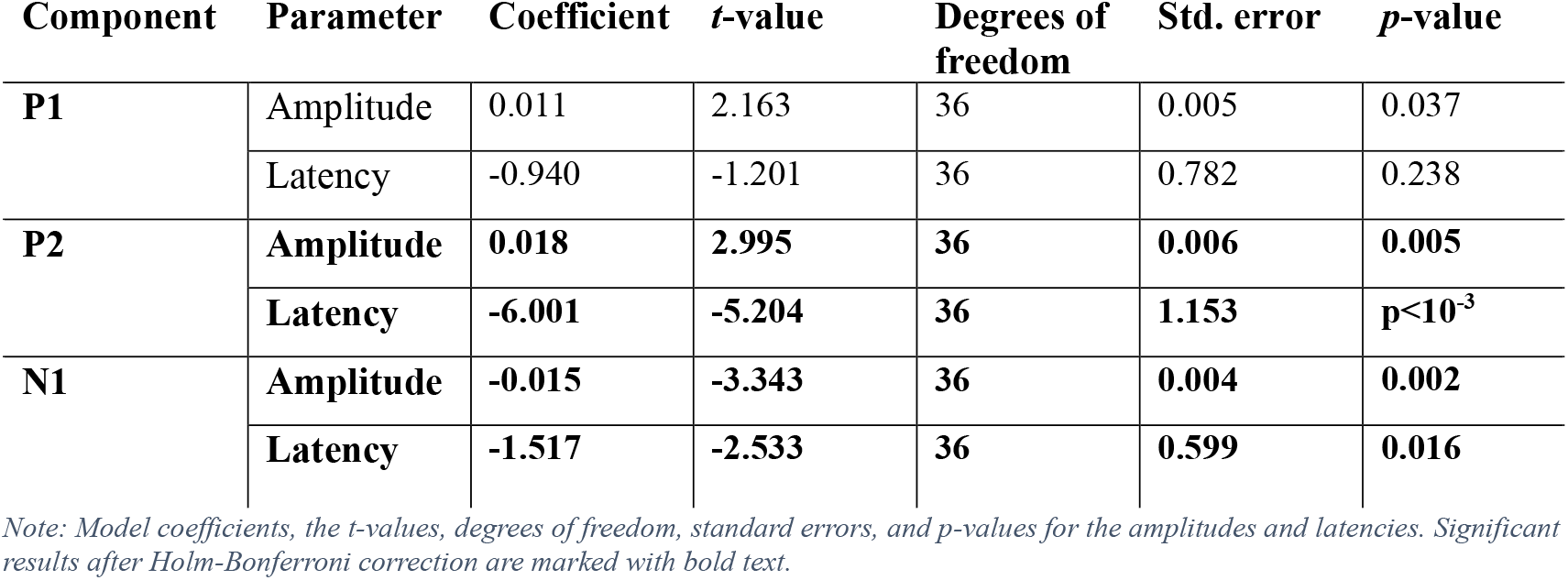
Statistical results from a linear mixed-effects model examining latencies and amplitudes.

**Figure 5.**
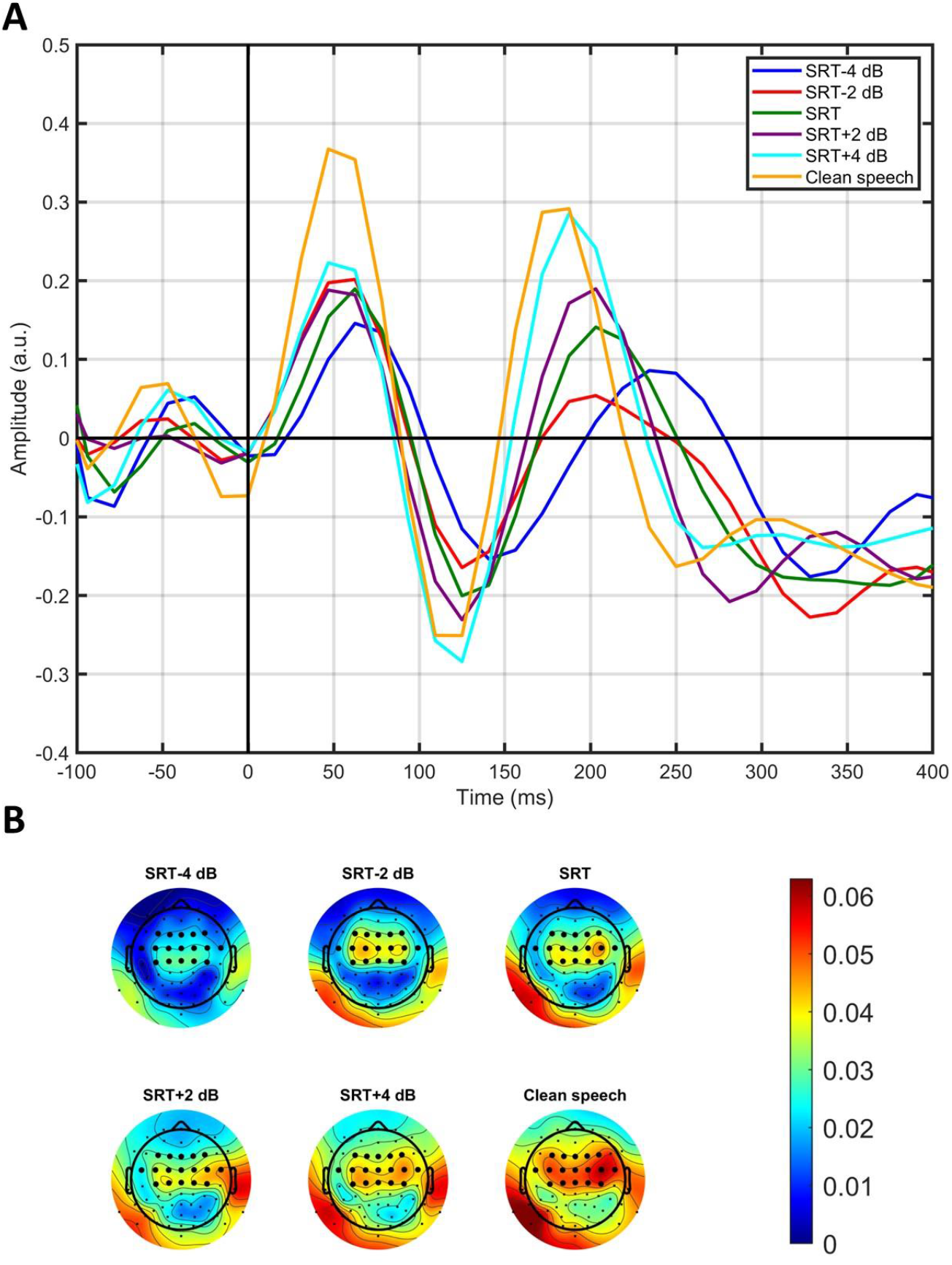
A: The grand average TRF in a -100 to 400 ms time window for each of the six SNR conditions, the last 100 ms of the time window is not depicted due to edging effects when approaching 500 ms. B: The grand average prediction accuracy from the encoding model; the electrodes used for the TRF analysis are shown in bold.

**Figure 6.**
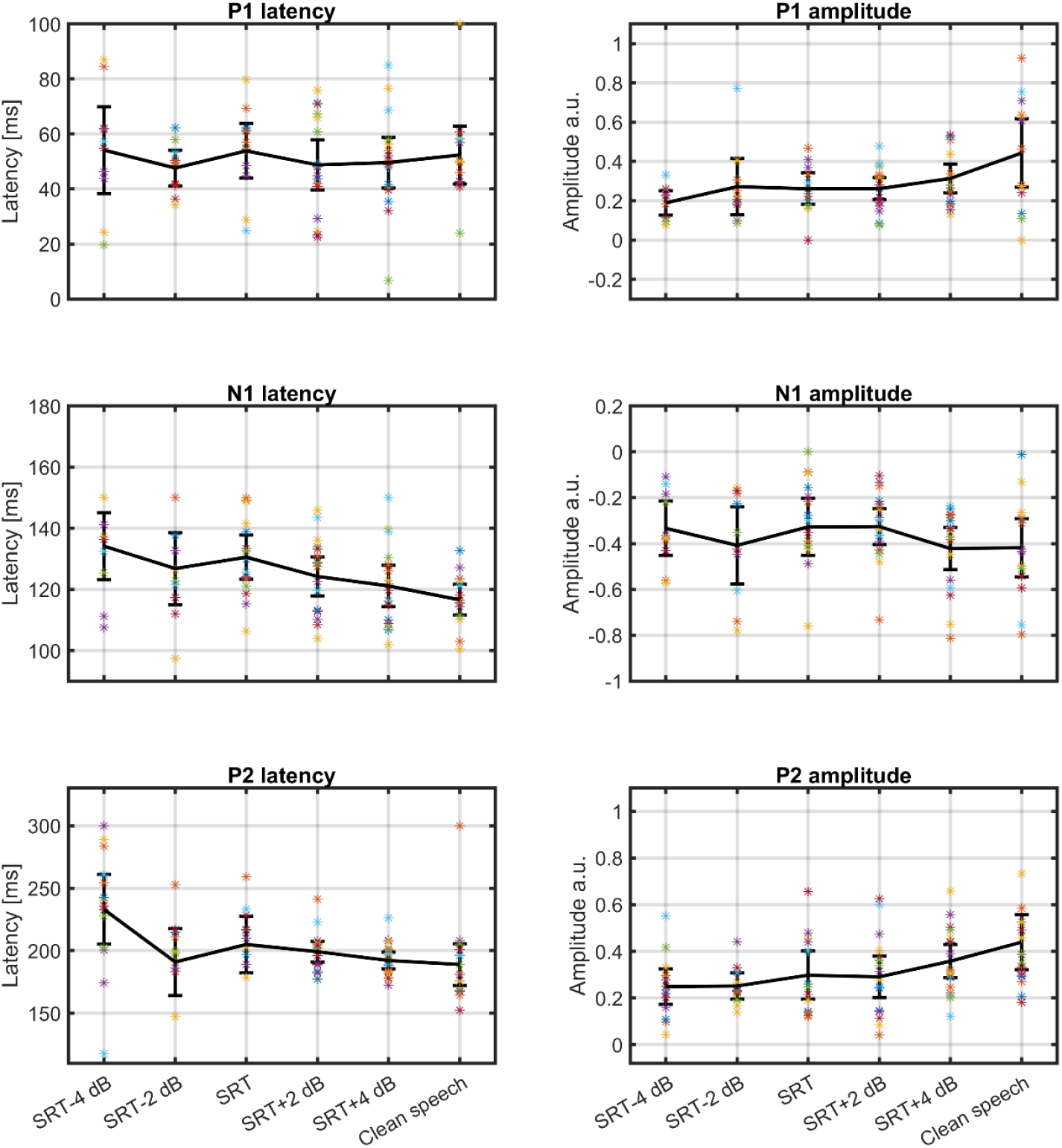
The across-participant average latencies and amplitudes of the TRF deflections are plotted along with the 95% confidence interval and the value for individual participants are depicted as colored stars.

## 4 Discussion

### 4.1 SRT estimation

The measured behavioral SRT (SRT_beh_) and the SRT estimated from the EEG data based on the reconstruction accuracy (SRT_neuro_) were almost identical on the group level, with median values of - 5.43 dB and -5.17 dB, respectively. When analyzing the results on the level of individual participants, we found that the SRT_neuro_ was within 2 dB from the SRT_beh_ for 15 out of 20 participants (75%), and within 3 dB from the SRT_beh_ for all participants (100%), as shown in Fig 4B. The maximum test-retest difference for the sigmoid fitting procedure within participant was 1.42 dB, indicating the robustness of the fitting procedure. The standard deviation of the difference between SRT_neuro_ and SRT_beh_ was 1.45 dB. It should be noted for reference that the average within-participant standard deviation for the Danish HINT is 0.86 dB for sentence-based scoring, resulting in the variation of the difference between test and retest of 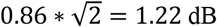 (Nielsen and Dau, 2011; Rønne et al., 2017). The standard deviation of the difference between SRT_neuro_ and SRT_beh_ is 1.45 dB and thus close to the test-retest variation expected for multiple measurements of the SRT_beh_ itself. However, while some of the variability between SRT_beh_ and SRT_neuro_ may be due to the limited precision of the reference measure SRT_beh_, part of it may also originate from other sources. For example, the reconstruction accuracy can be affected by attention (Ding and Simon, 2012; Mesgarani and Chang, 2012; O’Sullivan et al., 2015), and the quality of the EEG recordings can vary as well (Wilroth et al., 2023), contributing to the overall variation.

Prediction of SRT from EEG has been done in a study by Lesenfants et al. (2019), in which the SRT_neuro_ was estimated within 2 dB from the SRT_beh_ for 8 out of 19 participants (42%) using a delta-band envelope-based encoder. However, several differences exist between the two studies. The speech material to determine SRT_neuro_ differed substantially from the current study, with Lesenfants et al. (2019) using concatenated Flemish matrix sentences instead of the continuous-speech stimuli utilized in the current study, which could potentially affect participant engagement and attention fluctuations. If the test-retest standard deviation differed for the two speech tests used to determine the SRT_beh_, this could also explain a higher variance in one compared to the other. This does not appear to be the case here, as Lesenfants et al. found a within-participant standard deviation of 0.8 dB when using 6 noise conditions for determining the SRT_beh_, and 1.5 dB when using 4 noise conditions. These values closely align with the standard deviation for the Danish HINT (0.86 dB) (Lesenfants et al., 2019; Nielsen and Dau, 2011). In the current study, the SRT_beh_ values ranged from -6.00 to -3.28 dB (a 2.72-dB range) with a mean of -5.35 dB, while Lesenfants et al. (2019) observed a range of -10.3 to -4.7 (5.6-dB range) with a mean of -7.1 dB. The narrower range and higher mean SRT_beh_ in the current study can likely be attributed to the use of an open-set sentence test, whereas Lesenfants et al. used a matrix speech test for determining the SRT_beh_. To predict the envelope of the stimuli, Lesenfants et al. (2019) used a different model, an encoder, with a shorter time window (0 to 400 ms) compared to the current study (-100 to 350 ms). Furthermore, the EEG data in Lesenfants et al. (2019) underwent bandpass filtering with a narrower filter (0.5-4 Hz) than used in the current study (1-8 Hz). Moreover, Lesenfants et al. (2019) used a grand-average model to reduce experimental time. Per participant in Lesenfants et al. (2019), 14 min of EEG data were recorded with clean speech for model training and 6-8 min of EEG data were recorded for each noise condition, with either 4 or 6 noise conditions considered, depending on which group the participant got sorted into. Therefore, each participant in the Lesenfants et al. (2019) study had 2 min less EEG data collected with clean speech for model training and 8-10 min less test data per noise condition than in the current study.

The SNRs used in the noise conditions ranged from -12.5 to 2.5 dB in the Lesenfants et al. (2019) study, whereas the SNRs in the current study ranged from -10.00 to 0.72. Thus, the SNR levels are comparable across the two studies, despite the differences in SRTs. In the Lesenfants et al. (2019) study, the SNRs were constant across participants, in contrast to the current study, where the SNRs were chosen in an ordered manner in 2-dB steps around the SRT_beh_. To assess whether this selection of SNRs potentially biased the midpoint of the sigmoid fits toward the SRT_beh_, we applied the sigmoid fitting procedure to 5 pseudorandom reconstruction accuracies drawn from a standard uniform distribution between realistic flooring and ceiling values. We did not find that the SNR levels biased the SRT estimate, as multiple iterations of this random procedure did not yield meaningful SRT estimates. By centering the SNRs around the SRT_beh_, we increased the probability of measuring meaningful reconstruction accuracy values between flooring and ceiling of the sigmoid function. For a young normal-hearing population, this is not necessarily beneficial since the range of SRT_beh_ is narrow, which is also the case in the current study (see Fig 4B). However, if conducting a similar study with a population exhibiting a wider range of SRT_beh_, or e.g. evaluating a hearing-aid algorithm, this approach could prove advantageous, because the informative range is measured. On the other hand, the approach presumes that the behavioral SRT is available, which cannot necessarily be assumed, especially when considering meaningful use cases of the EEG-based SRT estimation. For instance, when measuring a non-responsive population, the behavioral SRT cannot be measured, and it may be necessary to sample a wider range of SNRs than in the current study to find informative levels.

The range of reconstruction accuracies of the decoder (expressed as Pearson’s correlations, see Fig. 4A) in the current study is comparable to those reported in previous studies (Lesenfants et al., 2019; Vanthornhout et al., 2018). A statistically significant increase in reconstruction accuracies was observed when increasing the SNR in 2-dB steps in the interval of *SNR* = SRT_beh_-4 dB to *SNR* = SRT_beh_+4 dB. This result indicates that the selected SNR range is relevant (with no flooring or ceiling effects) and that the 2-dB step size is appropriate, as also supported by visual inspection of the reconstruction accuracy-vs-SNR data points in Fig. 4A. However, the optimal step size may vary depending on the type of interfering signal and the characteristics of the population being tested, as both factors may influence the slope of the function.

It was not initially assumed that the SRT_neuro_ would be an unbiased estimate of SRT_beh_. However, no significant bias was observed, as the median of the individual difference between SRT_beh_ and SRT_neuro_ was very close to zero (0.38 dB). Consequently, no correction for a potential bias was applied. A bias correction could be necessary if changes in the acoustics or speech intelligibility measure were applied, for example if a different speech test was used for the SRT_beh_, or if different stimuli were used for SRT_neuro_ determination.

### 4.2 Temporal response function

The grand average TRF showed three distinct deflections within the time window from 0 to 400 ms, see Fig. 5A. When observing the latency and amplitude for these deflections (see Fig. 6), a general pattern emerged, with latencies decreasing and amplitudes increasing with increase in SNR. This pattern aligns with findings from a previous magnetoencephalography (MEG) study on normal hearing young adults (Ding and Simon, 2012) and an EEG study on preschool children (Van Hirtum et al., 2023b).

When using mixed models to investigate the effect of SNR on the latency and amplitude of the individual deflections across all noise conditions (excluding the clean speech condition due to the undefined SNR), a significant decrease in the latency of N1, P2 and a significant increase in the magnitude of the amplitude of the N1 and P2 were observed with increasing SNR (Table 3). However, interpretating the changes in the individual peaks of the TRF should be done with caution. Firstly, because the amplitude and latency of the TRF’s peaks and troughs are not uniquely tied to the amplitude and latency of the underlying neural components. For instance, a change in the amplitude of a single underlying component may impact not only the amplitude and latency of the local TRF deflection but also those of the neighboring deflections. Secondly, because the underlying neural components may be affected by preceding components. Still, the increased latencies with a lowering of the SNR could suggest that more neural processing is needed when the SNR is low.

The SNR-related changes in the TRF are interesting as they may be a result of the same changes in the underlying neural processes as the changes seen in the reconstruction accuracy. Specifically, a higher SNR results in a higher reconstruction accuracy, which may be attributed to more evident neural encoding of the stimuli, potentially expressed as higher amplitudes of the TRF.

Muncke et al. (2022) investigated the TRF responses at different SNRs in a normal-hearing population using concatenated German matrix sentences from the Oldenburg Sentence Test (OLSA) as stimuli. The interferer consisted of random overlapping OLSA sentences. Their findings showed a negative deflection around 100 ms and a positive deflection around 200 ms, with an increasing amplitude and decreasing latency with increasing SNR, consistent with the results of the current study. Additionally, the N1 amplitude in the TRF has been shown to increase and its latency decrease with higher amplitude of the stimuli when using amplitude binning of the envelope (Drennan and Lalor, 2019). This may also be part of the explanation for the increasing amplitudes and decreasing latencies observed with higher SNRs in this study, as modulation depth grows with increasing SNR. The previously observed increase in the decoders reconstruction accuracy with higher SNR is likely connected to changes in the TRF, were greater SNR – and thus larger modulation depth – leads to higher amplitudes and shorter latencies. After all, whether using a forward or a backwards approach, we are modelling the same underlying neural phenomena.

Notably, the prediction accuracy values in this study are higher centro-frontally compared to other scalp regions, and they increase with increasing SNRs, see Fig. 5B. Similar observations have been reported in other EEG studies using encoding models with speech stimuli (Drennan and Lalor, 2019; Fuglsang et al., 2017). We noticed that in the current study the TRF had not converged to zero at 400 ms. Hence, as a validation measure, we examined an encoder utilizing a longer time window (-100 to 1000 ms) to verify convergence towards zero, which was confirmed.

### 4.3 Limitations

The reconstruction accuracy of the envelope is not a direct measure of speech intelligibility. However, previous studies have shown that the cortical entrainment to the envelope of speech is very robust to background noise, and the auditory cortex synchronizes to the speech rather than the physical presented mixture of speech and noise (Ding and Simon, 2013). It has also been argued that the reconstruction accuracy of the envelope is correlated with speech intelligibility (Ding and Simon, 2013; Iotzov and Parra, 2019; Vanthornhout et al., 2018) and that the SRT_beh_ can be predicted from the reconstruction accuracy of the envelope (Lesenfants et al., 2019). A recent study (Karunathilake et al., 2023) suggests that the reconstruction accuracy of the envelope is not a direct speech intelligibility measure. In their study, vocoded speech was used as the stimulus, and speech intelligibility was manipulated by either presenting the clean speech version prior to presenting the vocoded speech or not. The results indicated that the envelope contributes significantly to the predictive power of an encoder, independent of the intelligibility. Nonetheless, speech intelligibility and envelope reconstruction are still correlated, likely because the access to envelope information is essential for speech understanding (Shannon et al., 1995). Lesenfants et al. (2019) observed an improvement in SRT_neuro_ estimation when using an encoder with both phonetic and spectral features. Further research is needed to determine if these features could also improve predictions in the current dataset.

In the study by Karunathilake et al. (2023), it was observed that there was no significant difference in TRF amplitudes when manipulating only the intelligibility of the speech stimuli while keeping the acoustics of the stimuli constant, using an encoder with envelope as a stimuli feature. However, significant differences in P1 and N1 amplitudes were found when comparing vocoded and non-vocoded speech which causes changes in the envelope of the speech signal, thus indicating that the changes we observed in TRFs may reflect envelope representation rather than a direct measure of speech intelligibility. In the Karunathilake et al. (2023) study, TRFs obtained from an encoder using word onsets showed two deflections, an early positive deflection (∼100 ms) and a later negative deflection (∼400 ms), both modulated by speech intelligibility, with a greater effect size observed for the later deflection. Further research is needed to determine if using word onsets instead of the speech envelope would provide a more direct measure of speech intelligibility.

The current study investigated a population of young normal-hearing participants, which results in a very narrow range of SRT_beh_ values and pure-tone thresholds, see Fig. 3. Further research is needed to evaluate the methods used in the current study with regard to individualized predictions in a more heterogeneous group exhibiting a wider range of SRT_beh_ values. Additionally, more research is needed to estimate the effect of different cognitive and auditory abilities on the TRF components and SRT_neuro_ estimation as the effect of cognitive and auditory abilities on the methods used in this study is unknown and previous studies have shown that cognitive and auditory abilities can influence speech comprehension (Dryden et al., 2017; Mukari et al., 2020; Nielsen and Dau, 2011; Ren et al., 2019).

## 5 Conclusion

Pearson’s correlation between the continuous speech envelope and the reconstructed speech envelope from EEG was used as a neural correlate of speech intelligibility. This was calculated at different SNR levels which was manipulated by controlling the level of steady state speech shaped noise. The EEG-based SRT (SRT_neuro_) was estimated by fitting a sigmoid function to the resulting reconstruction-accuracy-versus-SNR data, which was compared against the behavioral SRT (SRT_beh_). The SRT_beh_ and SRT_neuro_ matched almost perfectly on the group level without an obvious bias between the two SRT measures. For 15 out of 20 participants (75%), the SRT_neuro_ was within 2 dB of the SRT_beh_, and for all participants (100%) it was within 3 dB. In addition, exploration of the TRFs showed a significant decrease in N1 and P2 latency along with a significant increase of magnitude in N1 and P2 amplitude with increasing stimulus SNR.

## Abbreviations

(SRT_beh_): Behaviorally measured SRT
(EEG): Electroencephalography
(HINT): Hearing-In-Noise Test
(HL): Hearing level
(ICA): Independent component analysis
(LR): Likelihood ratio
(OLSA): Oldenburg Sentence Test
(SNR): Signal-to-noise ratio
(SRT): Speech-reception-threshold
(SRT_neuro_): SRT estimated from EEG
(SD): Standard deviation
(TRF): Temporal response function

## Data availability

The data used in the study is available upon reasonable request to the corresponding author.

## CrediT authorship contribution statement

**Heidi B. Borges:** Writing – original draft, Data curation, Formal analysis, Funding acquisition, Conceptualization, Software, Investigation, Visualization, Methodology. **Johannes Zaar:** Writing – review and editing, Funding acquisition, Conceptualization, Supervision, Methodology, Software, Project administration. **Emina Alickovic:** Writing – review and editing, Funding acquisition, Conceptualization, Supervision, Methodology, Software. **Christian B. Christensen:** Writing – review and editing, Conceptualization, Supervision, Methodology. **Preben Kidmose:** Conceptualization, Methodology, Writing – review and editing, Supervision, Project administration, Funding acquisition.

## Acknowledgements

The authors are thankful for all the participants who gave their time to be in this study. We would also like to thank Simon With for his support during the data collection. This work was supported by the William Demant Foundation [grant number 21-2912].

