## Appendix for "Speech-reception-threshold estimation via EEG-based continuous speech envelope reconstruction"

### Appendix
### A1: Temporal response functions for individual participants

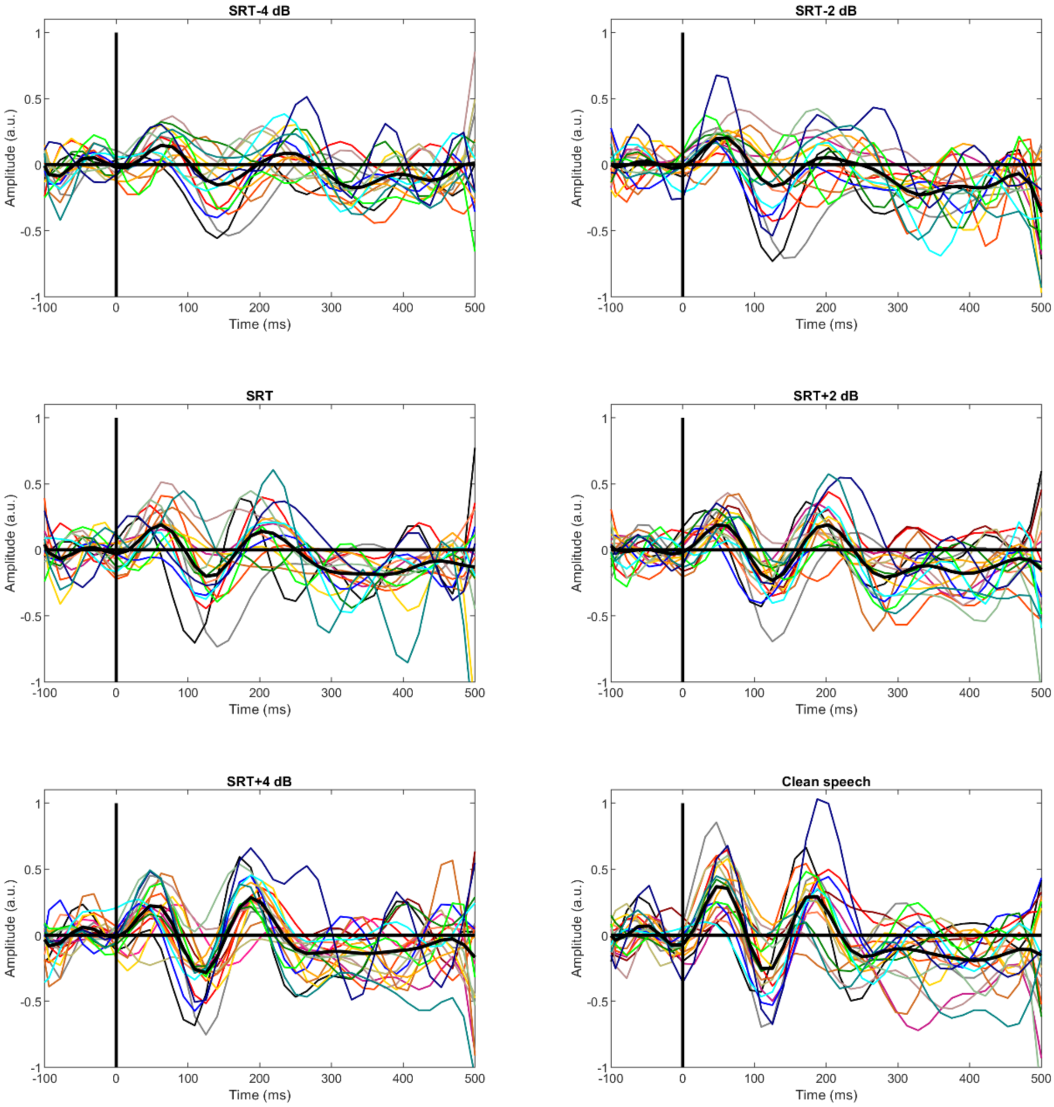


*1: The temporal response function for each participant and condition. Data for different participants are depicted as thin lines in different colors; the mean data across participants are shown as bold black lines. The conditions are evident from the figure titles.*
